# Differential Expression of the T cell Inhibitor TIGIT in Glioblastoma and Multiple Sclerosis

**DOI:** 10.1101/591131

**Authors:** Liliana E. Lucca, Benjamin A. Lerner, Danielle DeBartolo, Calvin Park, Gerald Ponath, Khadir Raddassi, David A. Hafler, David Pitt

**Author notes:** Correspondence to: David Pitt, MD, 300 George Street, Suite 353, New Haven, Connecticut 06511. These authors contributed equally to this work (Liliana E. Lucca, Benjamin A. Lerner).

## Abstract

To identify co-inhibitory immune pathways important in the brain, we hypothesized that comparison of T cells in lesions from patients with MS with tumor infiltrating T cells (TILs) from patients with GBM may reveal novel targets for immunotherapy. Focusing on PD-1 and TIGIT, we found that TIGIT and its ligand CD155 were highly expressed on GBM TILs but were near-absent in MS lesions, while lymphocytic expression of PD-1/PDL-1 was comparable. TIGIT was also upregulated in peripheral lymphocytes in GBM, suggesting recirculation of TILs. These data raise the possibility that anti-TIGIT therapy may be beneficial for patients with glioblastoma.

## Introduction

Immune checkpoint receptors are a family of co-inhibitory receptors that modulate T cell activation. The interactions between co-inhibitory receptors on tumor-infiltrating T cells and their ligands expressed by tumor cells is believed to contribute to the failure of the immune system to reject tumors [1,2]. Therapeutic blockade of this interaction has yielded dramatic results in the therapy of multiple cancer types. To date, this has led to FDA-approval of six immune checkpoint inhibitors that target cytotoxic T-lymphocyte-associated protein 4 (CTLA-4, Ipilimumab®) and programmed cell death protein 1 and its ligand (PD-1, Pembrolizumab® and Nivolumab®; PD-L1, Atezolizumab®, Avelumab® and Durvalumab®) [3–8].

To identify co-inhibitory pathways that may be important in the CNS, we hypothesized that comparison of T cells in lesions from patients with the autoimmune disease multiple sclerosis (MS) with tumor infiltrating T cells (TILs) of tumor from patients with glioblastoma multiforme (GBM) may reveal novel targets for immunotherapy in patients with CNS tumors. This approach was further suggested given the role in co-inhibitory and co-stimulatory pathways in T cell regulation that prevent activation of autoreactive T cells and that therapeutic blockade of checkpoint inhibitors is associated with a high incidence of autoimmune diseases ([9–11]). We focused on PD-1 and the T cell immunoreceptor with Ig and ITIM domains (TIGIT), a recently discovered co-inhibitory receptor expressed by activated T cells and natural killer cells [12,13]. TIGIT shapes T cell function by directly repressing pro-inflammatory Th1 and Th17 but not Th2 responses [14], and indirectly, by enhancing dendritic cell production of IL-10 [12]. TIGIT also prevents co-stimulatory signaling through CD226 by competing for the same ligand, CD155, and by disrupting CD226 homodimerization [15]. Finally, TIGIT is a marker of highly suppressive regulatory T cells (Tregs) and directly promotes Treg function in environments of Th1 inflammation [16, 17].

TIGIT has recently been proposed as a novel candidate target of cancer immunotherapy. Indeed, increased TIGIT has been demonstrated on tumor-infiltrating lymphocytes in a number of cancers including non-small cell lung cancer (NSCLC) and melanoma [15, 18]. Moreover, TIGIT blockade in animal models and in CD4 and CD8 T cells isolated from human tumors showed reinvigoration of anti-tumor immune responses [15, 19, 20]. However, TIGIT blockade also has the potential for inducing autoimmune disease, as expression of the competing co-stimulatory receptor, CD226, is increased on peripheral T cells of patients with rheumatoid arthritis and lupus [21]. Additionally, a coding variant in the CD226 gene is associated with multiple autoimmune diseases, including multiple sclerosis and rheumatoid arthritis [22]. Finally, TIGIT-deficient mice displayed increased susceptibility to developing experimental autoimmune encephalomyelitis (EAE), an animal model of multiple sclerosis [19,23], while treatment with a CD226-blocking monoclonal antibody delayed onset, and reduced severity of EAE [24].

We examined expression of TIGIT, CD226, their shared ligand CD155, and of PD-1 and its ligand PD-L1 in two prototypical neoplastic and autoimmune CNS diseases: glioblastoma and MS. Our data shows that TIGIT^+^ T cells were highly prevalent in glioblastoma infiltrates but not in in MS lesions, while the frequency of PD-1^+^ and PD-L1^+^ lymphocytes was comparable in the two conditions. Our findings highlight specific differences in immune checkpoint expression between glioblastoma and MS, and provide a strong rationale for developing immunotherapy against TIGIT for glioblastoma.

## Methods

### Tissue and blood samples

Immunohistochemistry was performed on formalin-fixed tissue from seven multiple sclerosis patients (obtained through autopsy) and seven glioblastoma patients (resection/biopsy). Flow cytometry was performed on blood from eight healthy volunteers, and on freshly resected glioblastoma tissue and matched blood from seven patients (one tumor did not yield enough T cells for analysis, Table 1). All samples were collected according to an Institutional Review Board-approved protocol; all patients and healthy volunteers gave informed consent.

**Table 1:**
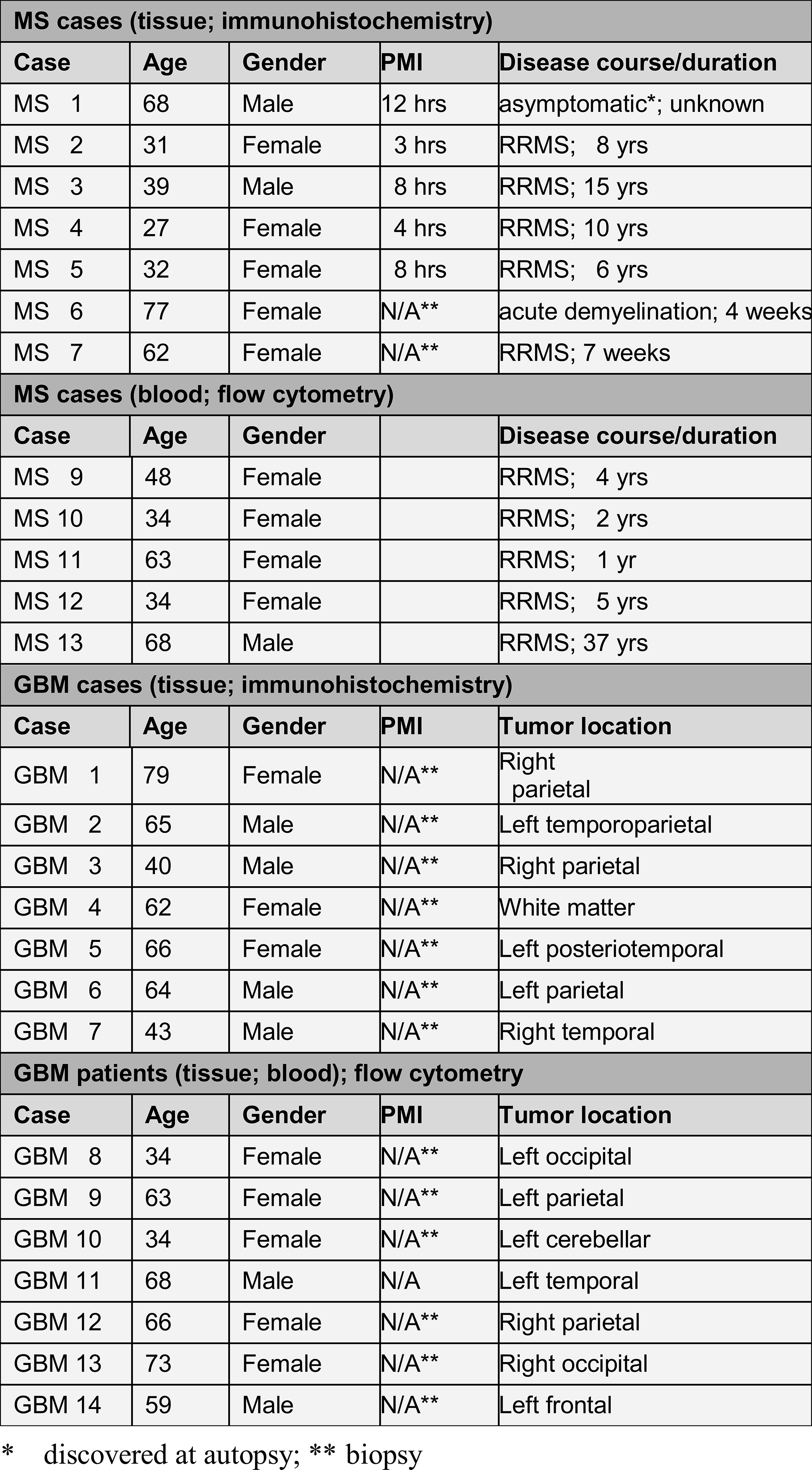
Clinical data of MS and GBM patients.

### Immunohistochemistry

Formalin-fixed paraffin-embedded sections were prepared for immunohistochemistry as described elsewhere [10]. Serial sections were stained with primary antibodies against CD3 (Dako A 40452, 1:200), TIGIT (Santa Cruz sc-103349, 1:800), CD226 (Novus Biologicals NBP1-85001, 1:50), and CD155 (Bioss Inc. bs-2525R, 1:750), and processed with the appropriate biotinylated secondary antibodies (Vector Laboratories, Burlingame CA) and avidin-biotin staining kit with diaminobenzidine as chromogen. Sections were counterstained with hematoxylin. Images were taken on a Leica DM5000 B microscope with Leica color camera DFC310 Fx Leica Application Suite (version 4.2.0) imaging software.

### Flow cytometry

Peripheral blood mononuclear cells were isolated from whole blood by Ficoll-Paque gradient centrifugation. Freshly resected tumor specimens were manually disrupted followed by digestion with collagenase IV (2.5mg/ml) and DNase I (0.2mg/ml) (Worthington Biochemical Corporation) for 1 hour. Tumor homogenates were separated on discontinuous 70-30% Percoll (Sigma Aldrich) gradients. Flow cytometric analysis was performed with antibodies targeting CD4 (BD Biosciences clone RPA-T4, V450 conjugated), CD8 (BD Biosciences clone RPA-T8, V500 conjugated), TIGIT (eBiosciences clone MBSA43, PerCP-eFluor ® 710 conjugated), CD226 (eBiosciences clone TX25, FITC conjugated), and PD-1 (BD biosciences clone EH12.1, Alexa Fluor ® 647 conjugated). Cell viability was assessed using Live/Dead Cell Viability Assays (Life technologies). Samples were run on a BD LSRFortessa or BD FACSAria II, as previously described. FlowJo software (Tree Star Inc.) was used for analysis after gating on live cells, with doublet exclusion followed by gating on CD4 and CD8 T cells.

### Statistics

Statistical analysis was performed in GraphPad Prism. P-values < 0.05 were considered significant.

## Results

We examined the expression of TIGIT, CD226, CD155, PD-1 and PD-L1 in tumor-infiltrating T cells in glioblastoma and demyelinating lesions from MS patients by immunohistochemistry. The number of TIGIT positive T cells was substantially higher in GBM infiltrates compared to MS lesions (Fig. 1A, C). Similarly, CD226 positive lymphocytes were more prevalent in glioblastoma than in MS, although cellular expression was overall low and had a punctate appearance. CD155, the ligand for TIGIT and CD226, was present at a low level in infiltrating lymphocytes in glioblastoma and MS, but was highly expressed by GBM tumor cells, perivascular monocytes and, to a lower degree, by cortical neurons (Fig. 1B). Finally, the number of infiltrating lymphocytes that expressed PD-1 and PD-L1 was similar in GBM and MS (Fig. 2A, B). In addition, we compared the number of TIGIT and CD226-expressing lymphocytes in perivascular cuffs and in the lesion parenchyma. In GBM, we found that the percentage of TIGIT^+^ T cells was significantly higher in tumor tissue than in perivascular infiltrates. Furthermore, we observed a gradient in the opposite direction for CD226^+^ lymphocytes, with higher frequencies in perivascular cuffs compared to tumor parenchyma. In contrast, the prevalence of TIGIT^+^ and CD226^+^ lymphocytes did not differ significantly between perivascular and parenchymal infiltrates in MS lesions (Fig. 2C).

**Figure 1.**
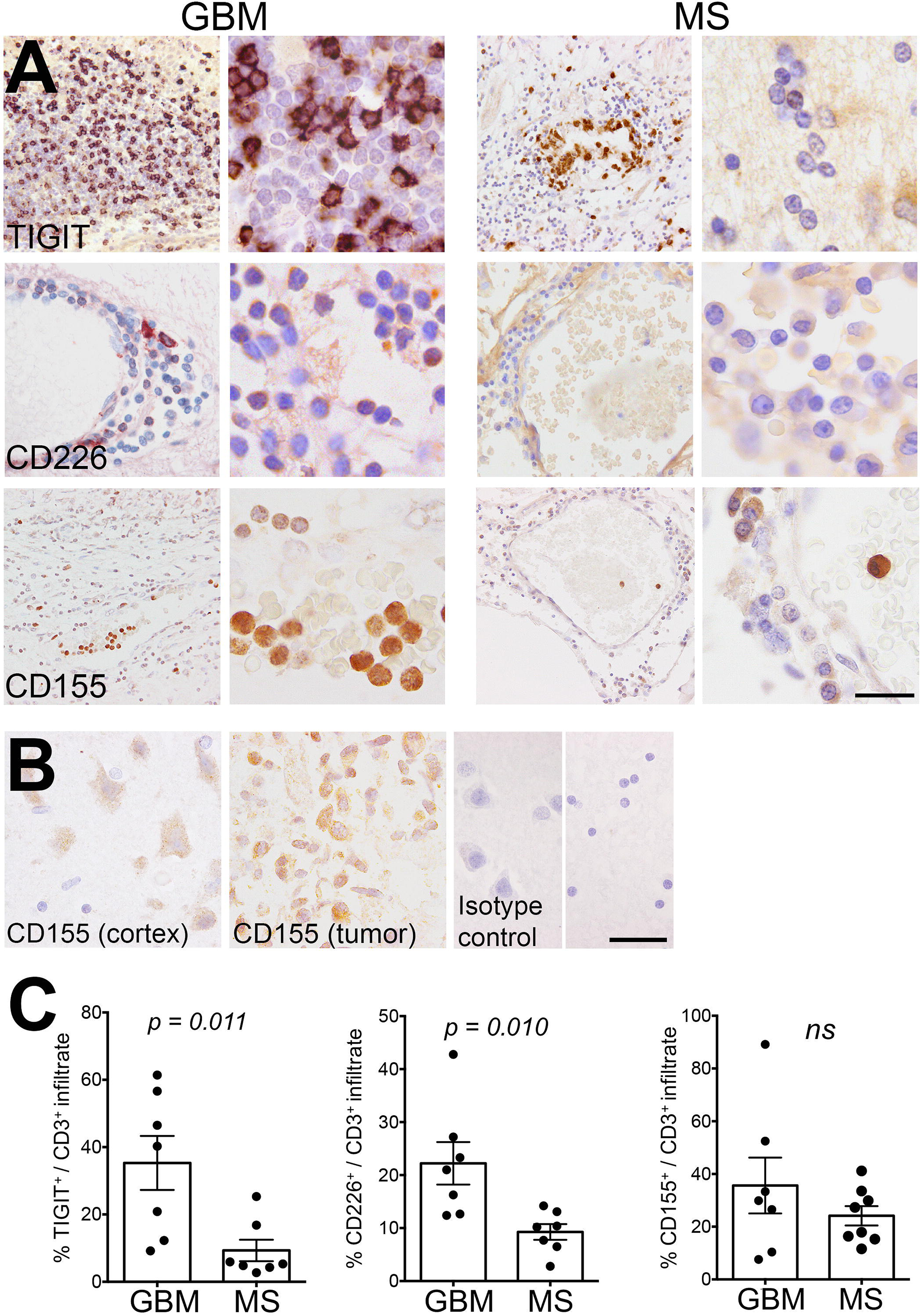
Expression of TIGIT and CD226 distinguishes GBM from MS T-cell infiltrates. (A) TIGIT^+^, CD226^+^ and CD155^+^ infiltrates in tumor tissue from GBM patients and chronic active lesions from MS patients. (B) CD155 expression in tumor cells and cortical neurons. (C) Quantification of TIGIT^+^, CD226^+^ and CD155^+^ infiltrates in GBM tumor tissue and MS lesions. Statistical significance was assessed by unpaired Student’s t tests with a p-value threshold of 0.05. ns = non-significant. Scale bars = 40 μm.

**Figure 2.**
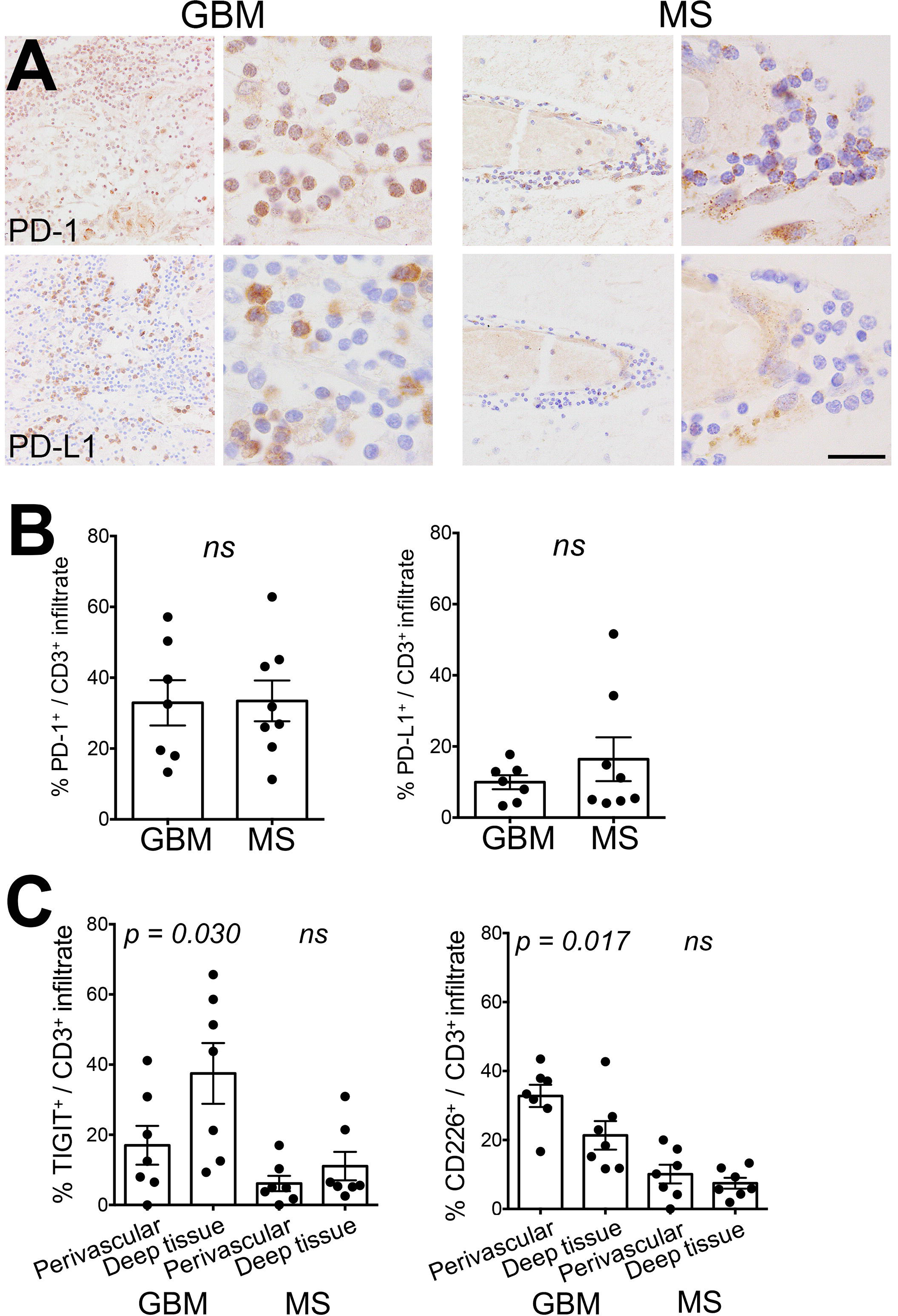
Expression of PD-1 and PD-L1 by T cell infiltrates is similar in GBM and MS. (A) PD-1^+^ and PD-L1^+^ infiltrates in tumor tissue from GBM patients and chronic active lesions from MS patients. (B) Quantification of PD-1^+^ and PD-L1^+^ infiltrates in GBM tumor tissue and MS lesions. (C) Frequency of TIGIT^+^ and CD226^+^ lymphocytes in the perivascular space and deep parenchyma. Statistical significance was assessed by unpaired Student’s t tests with a p-value threshold of 0.05. ns = not significant. Scale bar = 40 μm.

We used freshly resected (unfixed) tumor tissue from newly diagnosed glioblastoma patients to examine expression of TIGIT and CD226 by tumor-infiltrating T cells with flow cytometry. We observed that the majority of CD8 but not CD4 T cells were TIGIT positive, while CD226 was expressed in similar numbers by both lymphocytic subsets (Fig. 3A, B). Moreover, since TIGIT and CD226 have been reported to interact in cis, i.e. on the same cell, we assessed co-expression of the two molecules by flow cytometry, and found that the majority of tumor-infiltrating CD8 T cells expressing TIGIT were also positive for CD226 (Fig. 3C).

**Figure 3.**
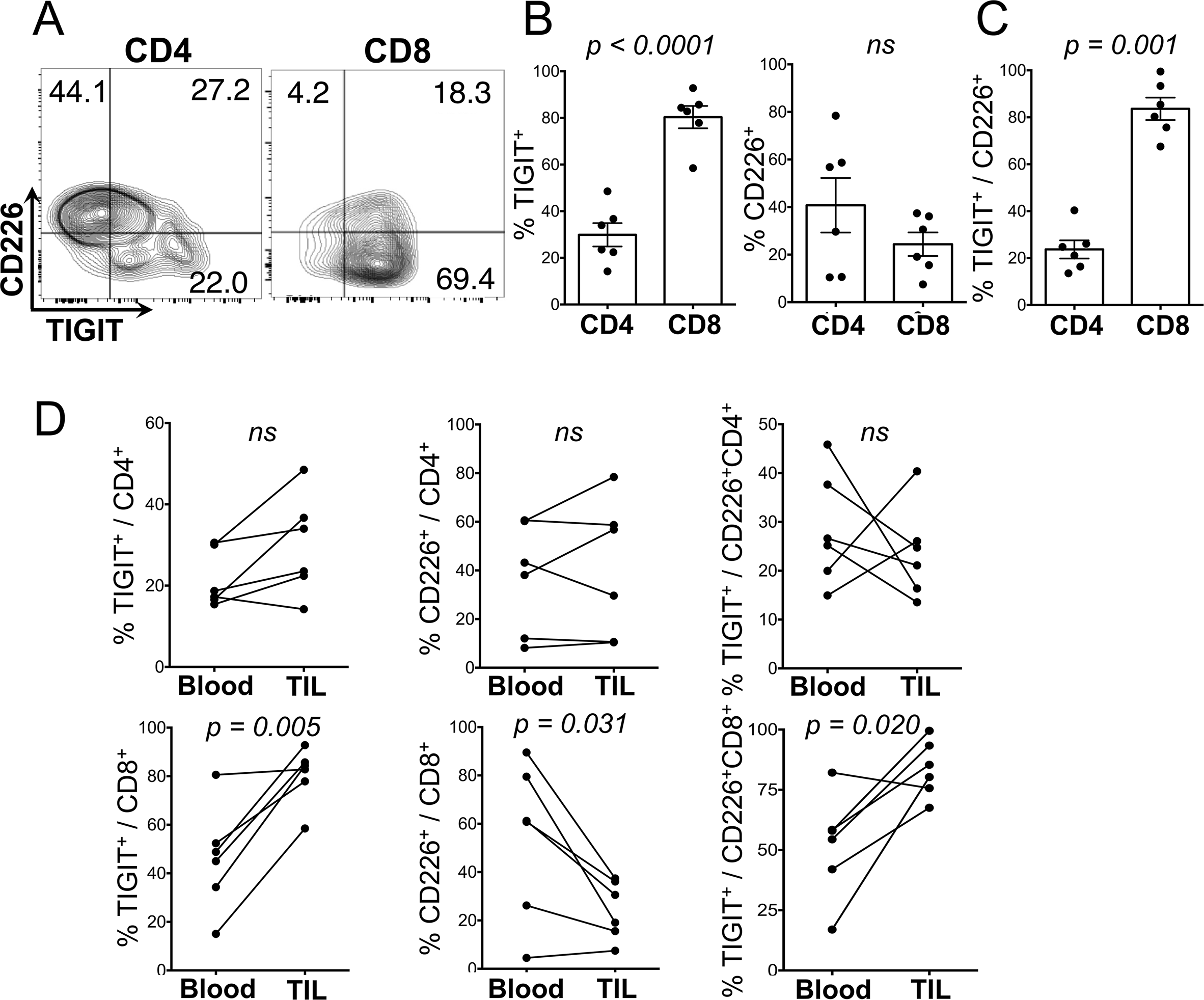
The tumor microenvironment of GBM allows for engagement of TIGIT on CD8^+^ T cells by CD155. (A) Expression of TIGIT and CD226 measured by flow cytometry on CD4 and CD8 T cells from tumor infiltrates. (B) Percent of CD4 and CD8 T cells expressing TIGIT, CD226, and percent of CD226^+^, CD4, and CD8 T cells co-expressing TIGIT (C). Histograms represent mean ± S.E.M. (D) Quantification of the frequency of TIGIT^+^, CD226^+^, and TIGIT^+^ among CD226^+^ for CD4 and CD8 circulating and tumor-infiltrating T cells. Frequencies were assessed by flow cytometry. Dots connected by a line represent the same patient. TIL = tumor infiltrating lymphocytes. Statistical significance was assessed by paired Student’s t test with a p-value threshold of 0.05; ns = non-significant.

Finally, we compared TIGIT/CD226 expression in tumor-infiltrating and blood-derived lymphocytes of glioblastoma patients and healthy controls. TIGIT^+^ CD8 but not CD4 T cells were increased in tumor tissue compared to peripheral blood of glioblastoma patients, while CD226^+^ CD8 lymphocytes were more prevalent in peripheral blood, where they co-expressed lower levels of TIGIT (Fig 3D). Overall, glioblastoma patients had a strong enrichment in TIGIT^+^ CD4 and CD8 T cells compared to healthy controls (Fig 4). The frequency of CD226^+^ CD4 and CD8 T cells and of PD-1^+^ CD4 and CD8 T cells was not significantly increased in the blood of GBM patients (Fig. 4C, D).

**Figure 4.**
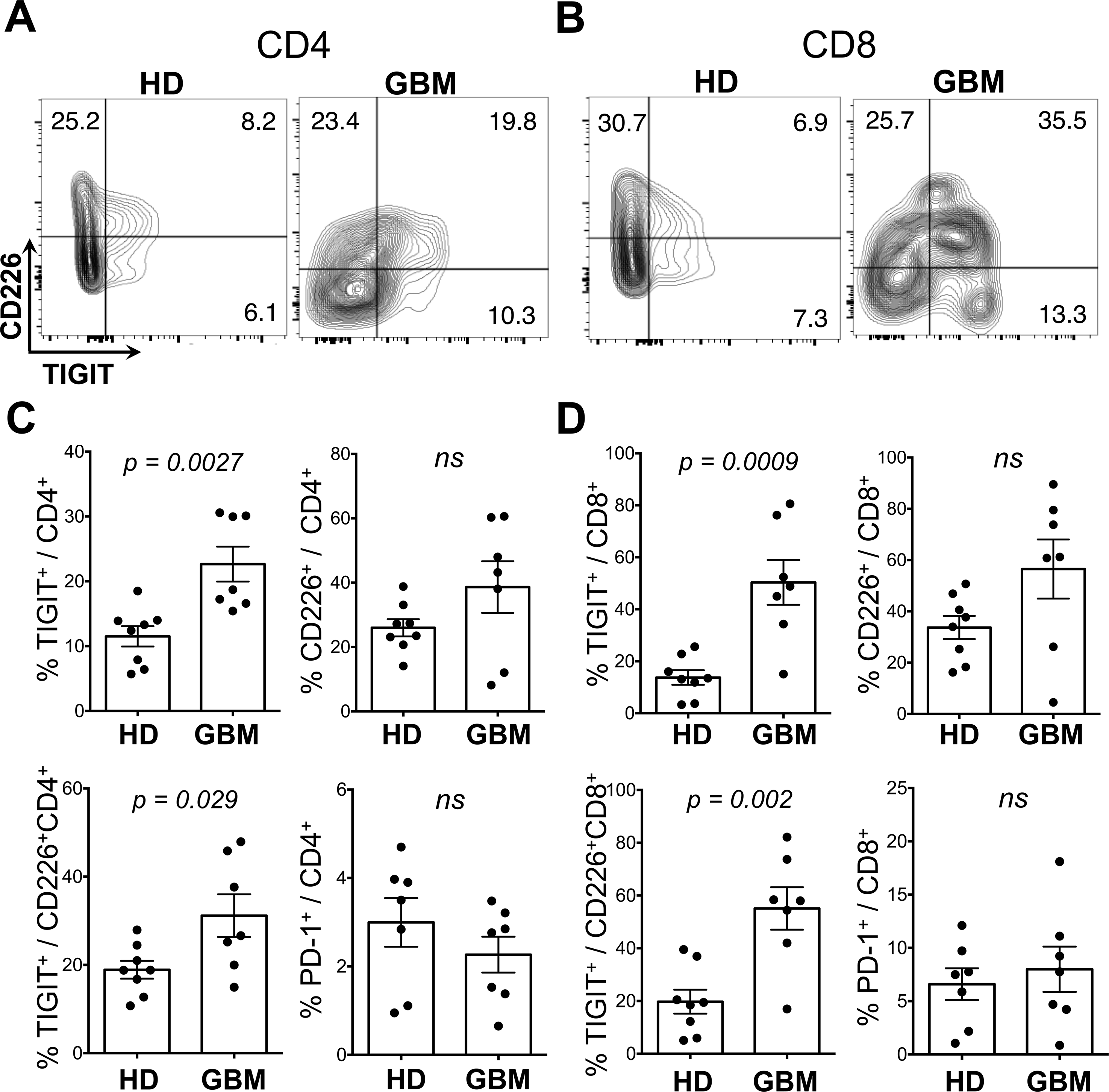
Circulating CD4 and CD8 T cells of GBM patients are enriched in TIGIT^+^ cells compared to healthy donors. Expression of TIGIT and CD226 measured by flow cytometry on circulating CD4 (A) and CD8 (B) T cells from healthy donors (HD) and GBM patients. Quantification of the frequency of TIGIT^+^, CD226^+^, and TIGIT^+^ among CD226^+^ and PD-1^+^ for CD4 (C) and CD8 (D) circulating T cells. The values for GBM are the same as depicted in Fig 2 in the “Blood” group. Histograms represent mean ± S.E.M. Statistical significance was assessed by unpaired Student’s t test with a p-value threshold of 0.05; ns = non-significant.

## Discussion

Immune checkpoint receptors are a family of co-inhibitory receptors that modulate T cell activation. The interactions between co-inhibitory receptors on tumor-infiltrating T cells and their ligands expressed by tumor cells is believed to contribute to the failure of the immune system to reject tumors [1,2]. While therapeutic blockade of this interaction has yielded dramatic results in the therapy of multiple cancer types, therapeutic trials with the immune checkpoint inhibitors anti-PD-1 and CTLA-4 in patients with glioblastoma have not been successful [25]. This suggests that PD-1 signaling might be redundant in the CNS, where other co-inhibitory pathways may be operative. To identify co-inhibitory pathways important in the brain, we hypothesized that comparison of T cells in lesions from patients with MS with TILs from patients with glioblastoma may reveal novel targets for immunotherapy in of brain tumors. Here we report that TIGIT expressing lymphocytes were substantially higher in glioblastoma infiltrates than in MS lesions. Previous studies have reported similar frequencies of TIGIT^+^ tumor-infiltrating lymphocytes in glioblastoma (25%-60%), although they have not been able to address whether those frequencies represent the level of TIGIT T cell expression in any CNS environment or a feature specific to the glioblastoma infiltrate [20, 26]. Given the abundant expression of the TIGIT ligand, CD155, on glioblastoma cells, this suggests that TIGIT signaling critically limits antitumor responses in GBM. In contrast, the relative absence of TIGIT/CD155 in normal white matter and MS lesions indicates that TIGIT signaling does not occur constitutively in the CNS. Moreover, PD1/PD-L1 positive lymphocytes were present in both conditions, indicating that PD-1 signaling is not a distinguishing feature between inflammatory responses in GBM and MS.

In tumor infiltrates, TIGIT was expressed by substantially more CD8 than CD4 cells, suggesting that TIGIT-mediated suppression of anti-tumor responses in glioblastoma affects primarily cytotoxic CD8 T cells, i.e. the lymphocyte subtype that directly interacts with tumor cells. This is in line with recent work in murine syngeneic CT26 and EMT6 tumor models, where TIGIT blockade enhanced CD8 T cell anti-tumor responses, but had little effect on CD4 T cells [15]. Although we observed a greater number of CD226-expressing lymphocytes in tumor infiltrates compared to MS lesions, its co-expression with TIGIT is likely to disrupt CD226 homodimerization [15] and thereby render CD226 non-functional. In contrast, the low expression rate of TIGIT in infiltrating lymphocytes in MS lesions suggests low TIGIT/CD226 co-expression, resulting in undisrupted CD226 function.

The increased percentage of TIGIT^+^ lymphocytes in tumor parenchyma compared to perivascular infiltrates, the high expression CD155 in glioblastoma cells, and the relative decrease of CD226^+^ T cells within the tumor tissue all suggest that CD155-induced TIGIT signaling is most pronounced in direct proximity to tumor tissue. Further investigations will elucidate whether the tumor microenvironment locally induces up-regulation of TIGIT and down-regulation of CD226, or preferentially attracts and retains TIGIT^+^CD226^-^ lymphocytes.

Finally, TIGIT expression was also increased in peripheral T cells in patients with glioblastoma compared to healthy controls, while expression of CD226 was decreased. This novel observation could indicate a systemic leakage of TIGIT-inducing factor and/or recirculation of TIGIT^+^ T cells between the periphery and the tumor bed. Trafficking of T cells between tumor and periphery would present an opportunity to gain insight into induction of TIGIT expression, to monitor the state of tumor-infiltrating lymphocytes, and to block TIGIT activity peripherally, thereby circumventing the obstacle of the blood-brain barrier for therapeutic antibodies.

In summary, our study provides evidence that the CD155/TIGIT axis plays a critical role in immune evasion of glioblastoma. This lays the basis for the development of immunotherapy that targets TIGIT signaling in GBM, either alone or in combination with anti PD-1 or anti-CTLA-4 checkpoint inhibition.

## Acknowledgements

We thank the operating room staff at Yale-New Haven Hospital and Jonathan Cruz for assistance in obtaining patient samples. This works was funded by National Multiple Sclerosis Society (RG 4866-A-2), the Gregory M. Kiez and Mehmet Kutman Foundation and National Institutes of Health Grants P01 AI045757 (DP) and NIH U19 AI089992, P01 AI073748, U24AI11867, R0AI22220, UM1HG009390, P01 AI039671, P50CA121974 the Nancy Taylor Foundation for Chronic Diseases, and grants NMSS grants RG-1802-30153CA and 1061-A-18 (DAH)

## Author contributions

B.A.L. and L.E.L. planned and performed experiments, analyzed data and wrote the manuscript. D.D., G.P., and V.K.P. performed experiments and analyzed data; K.R. provided technical support. D.A.H. and D.P. designed the study, planned experiments and wrote the manuscript.

